# The chromatin regulator HMGA1a undergoes phase separation in the nucleus

**DOI:** 10.1101/2021.10.14.464384

**Authors:** Hongjia Zhu, Masako Narita, Jerelle A. Joseph, Georg Krainer, William E. Arter, Ioana Olan, Kadi L. Saar, Niklas Ermann, Jorge R. Espinosa, Yi Shen, Masami Ando Kuri, Runzhang Qi, Timothy J. Welsh, Yufan Xu, Rosana Collepardo-Guevara, Masashi Narita, Tuomas P. J. Knowles

**Affiliations:** Centre for Misfolding Diseases, Yusuf Hamied Department of Chemistry, University of Cambridge, Cambridge, UK; Cancer Research UK Cambridge Institute, Li Ka Shing Centre, University of Cambridge, Cambridge, UK; Department of Genetics, University of Cambridge, Cambridge, UK; Cavendish Laboratory, Department of Physics, University of Cambridge, J J Thomson Avenue, Cambridge, UK; Yusuf Hamied Department of Chemistry, University of Cambridge, Cambridge, UK; Transition Bio Ltd., Maxwell Centre, JJ Thomson Avenue, Cambridge, UK; School of Chemical and Biomolecular Engineering, The University of Sydney, Sydney, Australia

## Abstract

The protein high mobility group A1 (HMGA1) is an important regulator of chromatin organization and function. However, the mechanisms by which it exerts its biological function are not fully understood. Here, we report that the HMGA isoform, HMGA1a, nucleates into foci that display liquid-like properties in the nucleus, and that the protein readily undergoes phase separation to form liquid condensates in vitro. By bringing together machine-leaning modelling, cellular and biophysical experiments and multiscale simulations, we demonstrate that phase separation of HMGA1a is critically promoted by protein–DNA interactions, and has the potential to be modulated by post-transcriptional effects such as phosphorylation. We further show that the intrinsically disordered C-terminal tail of HMGA1a significantly contributes to its phase separation through cation–π and electrostatic interactions. Our work sheds light on HMGA1 phase separation as an emergent biophysical factor in regulating chromatin structure.

## Introduction

Inside the nucleus of eukaryotic cells, chromosomal DNA is packed and highly organized in a structure known as chromatin.^1^ Chromatin organization is exquisitely modulated by the dynamic binding of a wide-range of architectural proteins.^2^ These include proteins such as linker histone H1 and members of the high-mobility group (HMG) superfamily.^3–5^

HMGs are among the most abundant and ubiquitous non-histone chromosomal proteins. They can be grouped in the HMGA, HMGB, and HMGN families. HMGs affect chromatin architecture by interacting with DNA, nucleosomes and/or other chromatin proteins. For example, HMGs compete with each other or other factors, such as linker histone H1, for chromatin binding sites.^6^ Such ability of HMGs to profoundly modulate chromatin structure is speculated to be intricately linked with many fundamental processes such as transcription activation/inhibition, DNA replication, DNA repair, integration of retroviruses into chromosomes.^7,8^

Within the HMG superfamily, the HMGA family proteins, including HMGA1 (with isoforms a and b) and HMGA2, are thought to be important players in fine-tuning chromatin organization and function. These proteins consist of three highly conserved DNA binding domains (‘AT hooks’, i.e., Pro-Arg-Gly-Arg-Pro). These AT hooks confer a higher affinity for binding to the minor groove of A/T-rich DNA sequences. HMGAs also contain a negatively charged C-terminal tail that is speculated to enable interactions with the positively charge histone tails within nucleosomes, and facilitate interactions with other proteins.^9^

Functionally, HMGAs have been shown to be highly expressed in the embryo and downregulated during differentiation,^10^ and their expression can be induced by mitogenic stimuli,^11^ which links HMGAs to cell proliferative events including cancer.^12,13^ Furthermore, HMGA expression levels have now also been linked to DNA damage response and oncogene-induced stress as well as senescence,^14,15^ and HMGA1 in particularly, was shown to be an essential component of senescence-associated heterochromatic foci (SAHFs).^15^

In addition to their role in global chromatin condensation, as seen in SAHFs, the prevalent view at a genetic level describes that HMGAs promote DNA accessibility by both decompacting chromatin and removing the steric barriers that nucleosome–nucleosome interactions may impose to transcription regulatory proteins such as RNA polymerase.^4^ However, mounting evidence now suggests that even within highly condensed constitutive heterochromatin regions, nucleosome interactions are more fluid and dynamical than previously postulated, and do not necessarily imply a steric barrier for dynamic chromatin modulators to the underlying DNA.^1,16–22^

Consistent with this liquid-like behaviour of nucleosomes, and in line with facile regulation, liquid–liquid phase separation (LLPS)^23–29^ of chromatin and its associated proteins has emerged as an important mechanism that may be responsible, at least in part, for the formation of intranuclear compartments, also termed nuclear condensate bodies.^2,30–32^ Within the LLPS framework for nuclear organization,^30,32,33^ multivalent proteins (including RNA-binding proteins and proteins with low complexity domains),^24,34–36^ RNAs^37–42^ and DNAs^32,33^ undergo a concentration-dependent demixing to yield biomolecular condensates.^43–45^ Accordingly, the formation of the nucleoli,^37,46,47^ nuclear speckles,^48^ PML bodies,^49^ and several other nuclear compartments that lack physical membranes have been attributed to LLPS. Furthermore, chromatin proteins found within the heterchromatic environment, like H1^50^ and the heterochromatin protein 1 (HP1),^2,32,33^ have been shown to phase separate in vitro and in cells.

Here, we report that the HMGA1 isoform HMGA1a can undergo LLPS in cell nuclei and form liquid droplets in vitro. Using machine learning tools, we predict HMGA1a’s propensity to phase separate from sequence-based analysis and demonstrate experimentally and by coarse-grained modelling that phase separation is facilitated by the presence of DNA. Molecular simulation suggests that HMGA1a phase separation is enhanced in the presence of DNA due to dominant Arginine-phosphate interactions and that it might be promoted by HMGA phosphorylation. In biophysical experiments, leveraging our PhaseScan high-resolution droplet microfluidics platform, we map the phase diagrams of recombinant human HMGA1a for a range of protein and DNA concentrations. In cell experiments, we find that HMGA1a nucleates into foci that display liquid-like properties within the nucleus of fibroblasts and cancer cells. These findings shed light on HMGAs phase separation as an emergent biophysical factor in regulating chromatin structure, and further highlight phase separation as a likely critical factor for nuclear chromatin organization.

## Results and discussion

### Machine learning analysis predicts HMGA1 to have a propensity to undergo DNA-mediated phase separation

We first examined the propensity of HMGA1a to undergo phase separation using a machine learning approach to identify LLPS-prone regions within the sequence and identify the driving forces behind this process.^53^ Specifically, we used a model trained on a dataset describing homotypic phase separation of a series of peptides and proteins. This analysis shows that HMGA is more prone to undergo phase separation in a homotypic environment than 6% of the human proteome, indicating that the protein is likely to undergo phase separation, albeit potentially less readily (e.g., at higher concentrations) than many LLPS-prone scaffold proteins. Also, an amino-acid-window based analysis search yielded regions with a score above 0.5 **(Figure 1a, top panel)**, suggesting that HMGA1a exhibits locally highly LLPS-prone regions.

**Figure 1.**
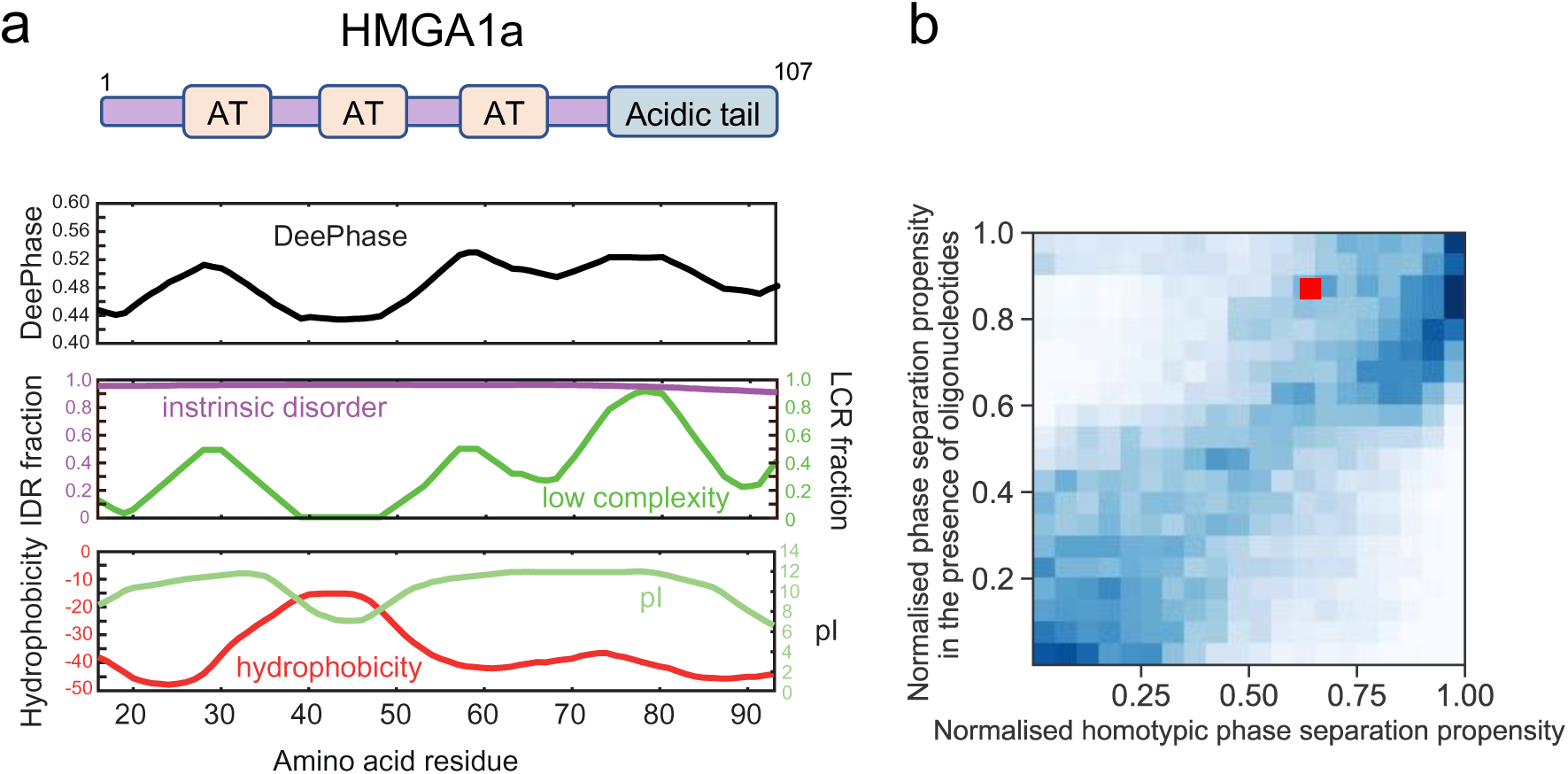
HMGA1a is a chromatin regulator protein that is predicted to undergo phase separation. **(a)** DeePhase phase separation score of HMGA1a and predictions of intrinsically disordered regions (IDR), low complexity regions (LCR), hydrophobicity, and isoelectric point (pI). The domain structure of HMGA1a is shown on top. AT hooks are denoted as AT. **(b)** Comparison of the phase separation propensity of HMGA1a with the human proteome (20,300 proteins) under homotypic conditions (x-axis) and in the presence of oligonucleotides (y-axis). HMGA1 noticeably moves up in the distribution, suggesting that the presence of oligonucleotides would promote its phase separation.

We further examined how some of the key physical parameters that are likely to govern phase separation vary across the sequence.^51,52^ Proteins that undergo phase separation in vitro and in vivo commonly contain intrinsically disordered regions that are marked by low sequence complexity.^34,35,49,53–58^ These features enable proteins to establish multivalent homotypic and/or heterotypic interactions with their binding partners necessary to drive phase separation.^35,36,43,59^ In agreement with this idea, analysis of HMGA1a for these features revealed that the protein is predicted to be intrinsically disordered **(Figure 1a, centre panel)** and exhibits regions within the protein that are of low sequence complexity **(Figure 1a, centre panel)**. These regions are rich in polar residues, as also indicated by a negative hydrophobicity score **(Figure 1a, bottom panel)**.

Notably, we also found that the isoelectric point (pI value) of the protein in these regions is high **(Figure 1a, bottom panel)**, corresponding to a net positive charge under physiological conditions. This trend suggests that phase separation propensity of HMGA1a may be enhanced further by the inclusion of negatively charged molecules, such as DNA. To probe this effect in more detail, we reparametrized our machine learning model to estimate additionally the phase separation propensities under conditions that included oligonucleotides **(Figure 1b)**. When comparing the score for phase separation in an heterotypic environment obtained from this latter model to the homotypic phase separation score for all the proteins across the full human proteome (20,300 proteins), we found that HMGA1a had moved from the 35th percentile under homotypic conditions to around the 12th percentile in an heterotypic environment **(Figure 1b)**, suggesting that the presence of oligonucleotides can notably enhance the phase separation of HMGA1.

Taken together, our sequence analysis suggests that HMGA1a has a propensity to undergo phase separation and that this process could be driven by the highly disordered nature of the sequence and the presence of low complexity regions that have high content of polar amino acid residues as indicated by a low hydrophobicity value. Furthermore, the analysis reveals a propensity to undergo LLPS is enhanced in the presence of DNA. Interestingly, similarly charged proteins, such as the disordered histone tails and the C-terminus of the linker histones, phase-separate in the presence of DNA and nucleosomes.^30,50^ Hence, the net positive charge of HMGA1a may serve to target DNA regions but also may contribute to its phase separation propensity.

### HMGA1a phase separation is driven by DNA in silico

To further investigate the ability of HMGA1a to phase separate and to gain a more detailed molecular understanding of the process, we used molecular dynamics simulations to examine HMGA1a LLPS in silico. To this end, we performed direct coexistence simulations using the hydrophobic scale (HPS) coarse-grained model of Dignon and colleagues^58^ with a modification of Das and colleagues^60^ that enhances cation–π interactions.

First, we conducted simulations on 48 copies of interacting full-length HMGA1a proteins (107aa). These simulations indicate that wildtype HMGA1a is unlikely to undergo phase separation without the aid of additional molecules or modifications **(Figure 2a (black binodal) and Figure 2b (bottom panel))**; within our energy scale, phase separation was only observed at very low temperatures (i.e., < 200 K model temperature). This result suggests that HMGA1a may require crowders or very high protein concentrations to undergo phase separation in vitro. Our simulations also reveal that the weak homotypic self-interactions among HMGA1a proteins involve mainly the C-terminal portion **(Figure 2c)**, which is in accordance with DeePhase and sequence-based analysis results above.

**Figure 2.**
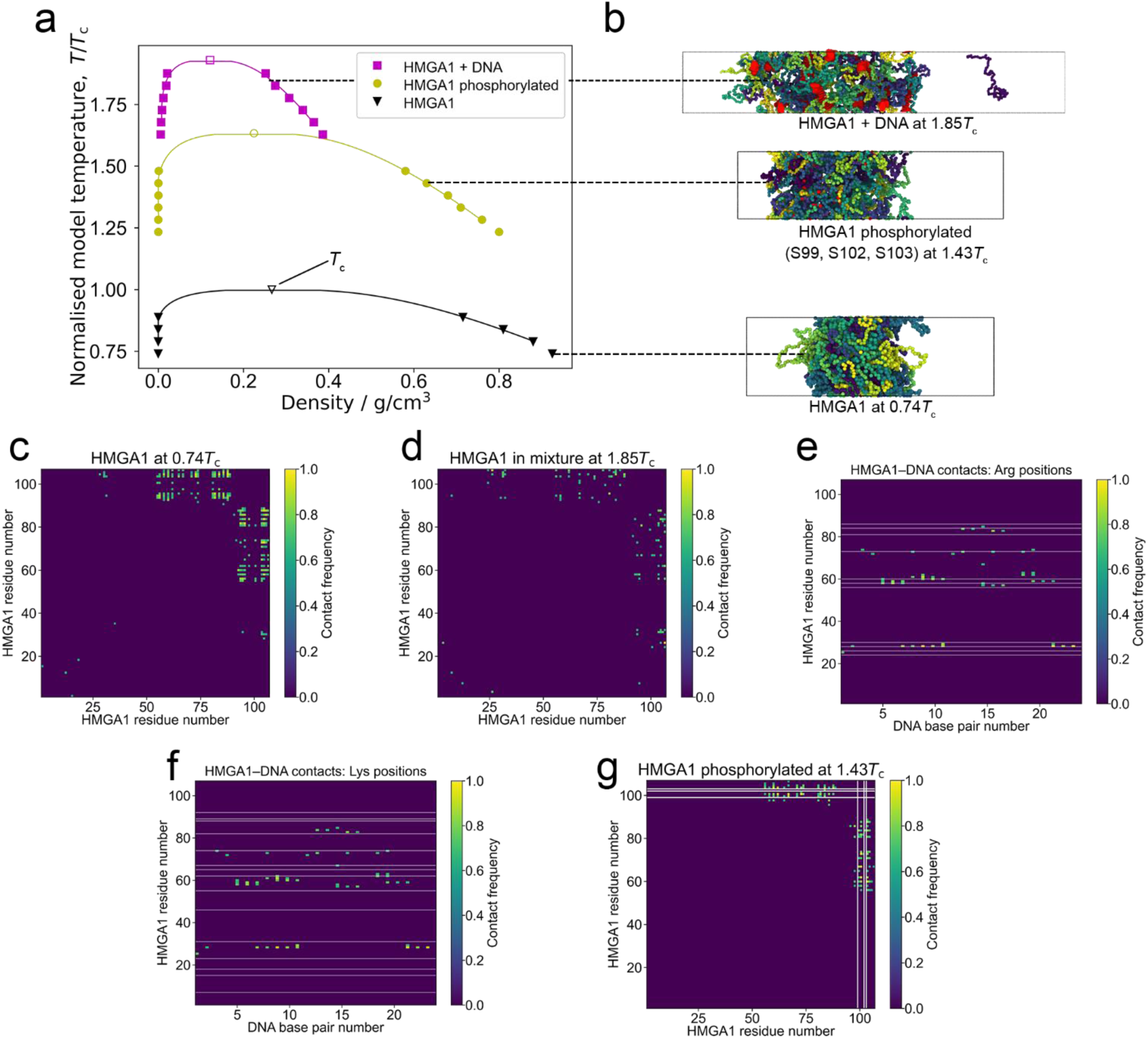
Molecular modelling suggests that DNA promotes phase separation of HMGA1a. **(a)** Phase diagrams (temperature versus density) for the wildtype (wt) HMGA1a protein (black triangles), the phosphorylated HMGA1a protein (yellow spheres), and the HMGA1a–DNA mixture (magenta squares). Estimated critical points (empty symbols) are given for each data set. Each binodal is normalised based on the critical temperature of the wildtype protein (*T*_c_ (wt)). **(b)** Snapshots from Direct Coexistence simulations of HMGA1a in an elongated box at 0.74 *T*_c_ (wt) (48 chains; bottom panel), of phosphorylated HMGA1a at 1.43 *T*_c_ (wt) (48 chains; middle panel), and the HMGA1a–DNA mixture at 1.85 *T*_c_ (wt) (48 protein chains + 12 DNA strands; top panel). **(c,d)** Amino acid contact maps for HMGA1a homotypic interactions in the (c) pure wildtype system at 0.74 *T*_c_ (wt), and (d) with DNA present at 1.85*T*_c_. Residues near C-terminal make most significant contributions to protein–protein interactions. **(e,f)** Contact map between HMGA1a residues and DNA base pairs with (e) arginine and (f) lysine residues indicated as horizontal white lines. Regions of high contact mostly coincide with positions of arginine residues. **(g)** Amino acid contact map for HMGA1a homotypic interactions in the phosphorylated system at 1.43 *T*_c_ (wt); positions of phosphorylation (S99, S102, and S103) are indicated as white lines. For clarity, in contact maps (c)–(g), all contact frequencies <0.6 are set to zero. Please see further details on calculations in Methods.

In a next set of simulations, we added short strands of double-stranded DNA (each 24 bp, consistent with average DNA linker lengths) to our solution of HMGA1a proteins. Here DNA is modelled using our chemically accurate coarse-grained model that captures the sequence and mechanical properties of DNA.^22^ DNA significantly increases the critical temperature of HMGA1a (i.e., >*1.9 T*_*c*_(wt)) indicating that LLPS of the unmodified HMGA1a protein is likely DNA-dependent **(Figure 2a (magenta binodal), Figure 2b (top panel))**. We also assessed the contact frequencies between HMGA1a molecules in the HMGA1+DNA mixture **(Figure 2d)** and between HMGA1a residues and the DNA strands **(Figure 2e,f)**. We found that HMGA1a molecules interact with each other in a similar manner to the wildtype system (i.e., predominantly via their C-tails). In terms of interactions with DNA, the contact pattern closely matches the patterning of Arg residues in the protein **(Figure 2e)** (see the comparison with Lys residues, **Figure 2f**). These results suggest that cation–π and electrostatic interactions between HMGA1a and DNA could drive LLPS. We also estimated the valency of DNA in terms of the number of individual proteins that each strand can recruit. We find that, on average, 12 DNA base pairs can recruit about 2 HMGA1a proteins. Based on these results, we postulated HMGA1a proteins can act as a glue, bridging DNA within HMGA1a liquid condensates.

Inside cells, HMGA1a is highly regulated by the presence of post-translational modifications. In fact, HMGA1a is one of the most heavily phosphorylated proteins inside the nucleus,^61^ and consistent with its LLPS propensity, phosphorylation increases the residence time of HMGA1a within heterochromatin regions.^5^ We therefore hypothesized that a crucial feature modulating the phase behaviour of HMGA1a proteins in vivo may be phosphorylation. To investigate this, we phosphorylated Ser99, Ser102, and Ser103 in our simulations, which are located at the negatively charged C-terminus and are phosphorylated in vivo by the Casein kinase (CK2),^62^ and repeated our Direct Coexistence simulations of HMGA1a proteins (in the absence of DNA). Strikingly, our simulation shows that phosphorylation dramatically enhances the ability of HMGA1a to undergo phase separation without the aid of additional binding partners **(Figure 2a (yellow binodal), Figure 2b (centre panel))**. Consistently, phosphorylation amplifies C-tail–C-tail interactions in the protein **(Figure 2g)**. Hence, we speculate that the HMGA1a condensates we observed in the nucleus in regions of very low DNA concentration (vide infra), might be composed of phosphorylated HMGA1a proteins. Interestingly, serine phosphorylation has been shown to reduce the binding affinity of HMGs to DNA^63^ by 3-fold, which would explain the preferential exclusion of DNA from heavily phosphorylated HMGA1a condensates.

### Recombinant human HMGA1a undergoes phase separation in vitro

Next, we explored whether HMGA1a undergoes phase separation and forms liquid-like assemblies in vitro. To this end, we expressed and purified human HMGA1a protein and probed its phase behaviour. At room temperature, aqueous solutions of HMGA1a at 10 µM concentration (labelled with Alexa 647 at sub-stoichiometric amounts) spontaneously demixed at physiological salt concentration to form liquid droplets of ca. 1–2 μm in diameter in the presence of 5% polyethylene glycol (PEG) **(Figure 3a, left panel)**. Over time, HMGA1a droplets coalesced to form larger droplets of about 3 μm in diameter, corroborating their liquid-like character. Notably, in the absence of PEG, HMGA1a phase separated only at high concentrations, and only at the air–water interface where evaporation occurs **(Figure S1a)**. The requirement of high HMGA1a concentrations and PEG for HMGA1a LLPS is consistent with our simulation studies, which suggest that the pure wildtype protein is unlikely to phase separate on its own. Further characterisation of the biophysical properties of HMGA1a condensates in the presence of PEG, including their surface tension and viscosity is provided in the Supplementary Information **(Figure S1b, Supplementary Results)**.

**Figure 3.**
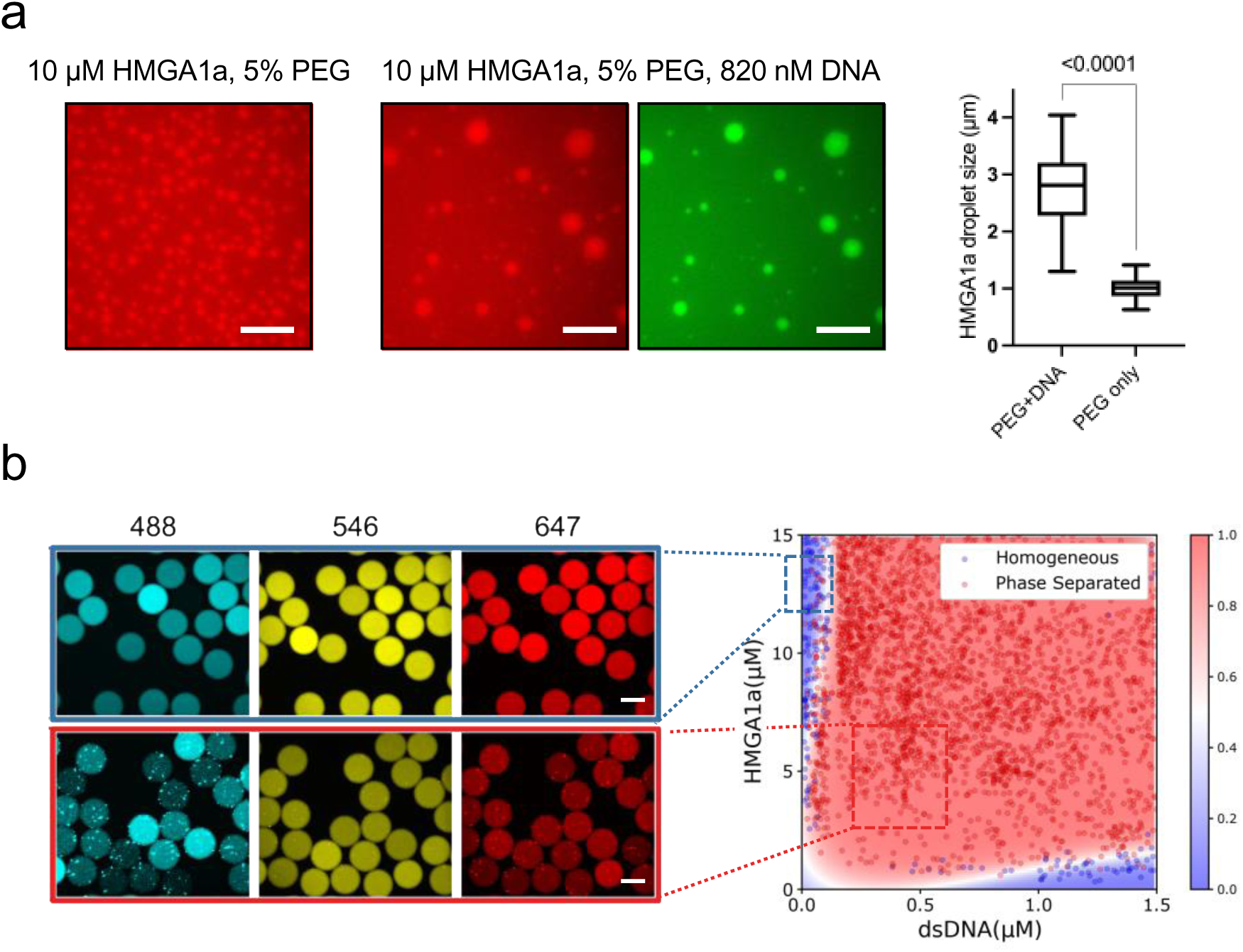
HMGA1a undergoes LLPS in vitro in the presence of crowding and DNA. **(a)** HMGA1a at 10 µM (labelled with Alexa 647, red) readily undergoes LLPS with 5% PEG (20k) (left panel) and 820 nM DNA (24 bp duplex, labelled with Atto488, green) present (centre panel). The comparison of HMGA1a condensate size with or without addition of DNA (right panel). Buffer: 50 mM Tris-HCl, 150 mM KCl. Scale bars, 10 μm. In the box plots, boxes extend from the 25th to 75th percentiles, with a line at the median. Whiskers span 1.5× the interquartile range. Statistical analysis was performed using a two-sided *t*-test (*p*-value < 0.0001). **(b)** Left panel: Epifluorescence microscopy images of microdroplets with Alexa 488 (blue), Alexa 546 (yellow), and Alexa 647 (red) fluorescence corresponding to DNA (24 bp duplex, labelled with Atto488), PEG, and HMGA1a (labelled with Alexa 647), respectively. Scale bars, 100 μm. Buffer: 50 mM Tris-HCl (pH 7.4), 120 mM KCl. PEG (20k) concentration was 3% (*w*/*v*). Right panel: Phase diagram of HMGA1a in the presence of DNA generated through PhaseScan droplet microfluidics. Red, phase separated, Blue, mixed. Phase diagram was generated from *N* = 6296. Colour bar: Probability of a region in the chemical space being phase separated classified and predicted using machine learning.

Given HMGA1’s role in modulating chromatin structure and the results of the sequence analysis and molecular simulations, we further probed HMGA1a phase separation in the presence of DNA. Aqueous solutions of HMGA1a at 10 µM concentration (labelled with Alexa 647) in the presence of nanomolar amounts of double-stranded DNA (labelled with Atto 488) readily formed condensates **(Figure 3a, centre panels)**. These condensates were up to 10-fold larger in size as compared to condensates in the absence of DNA under otherwise identical conditions **(Figure 3a, right panel)**. A control experiment without protein, but only DNA present, did not result in condensate formation **(Figure S2, right panel)**. This suggests that DNA promotes phase separation of HMGA1a in vitro.

To further characterise the phase behaviour of HMGA1a in the presence of DNA, we mapped out its phase diagram under constant crowding conditions. Using our PhaseScan high-resolution droplet microfluidics approach,^64^ we obtained phase diagrams for a range of HMGA1a and DNA concentrations **(Figure 3b)**. HMGA1a readily phase separated with minimal (i.e., nanomolar) amounts of DNA over a broad range of protein concentrations. Of note, excess of DNA or protein leads to dissolution of condensates. Moreover, droplets merged and coalesced, substantiating their liquid-like behaviour **(Figure S3)**.

### HMGA1a forms liquid droplet-like foci in the nucleus

Based on our in vitro findings, we then sought to examine whether HMGA1a forms phase separated condensates in cells. When EGFP-tagged HMGA1a was overexpressed under the strong CMV promoter in IMR90 human fibroblasts, we indeed observed the formation of droplet-like HMGA1a foci in the nucleus (**Figure 4a**). A closer look at the foci revealed that HMGA1a foci are spherical, consistent with liquid-like systems typically associated with LLPS.^59,65–67^

**Figure 4.**
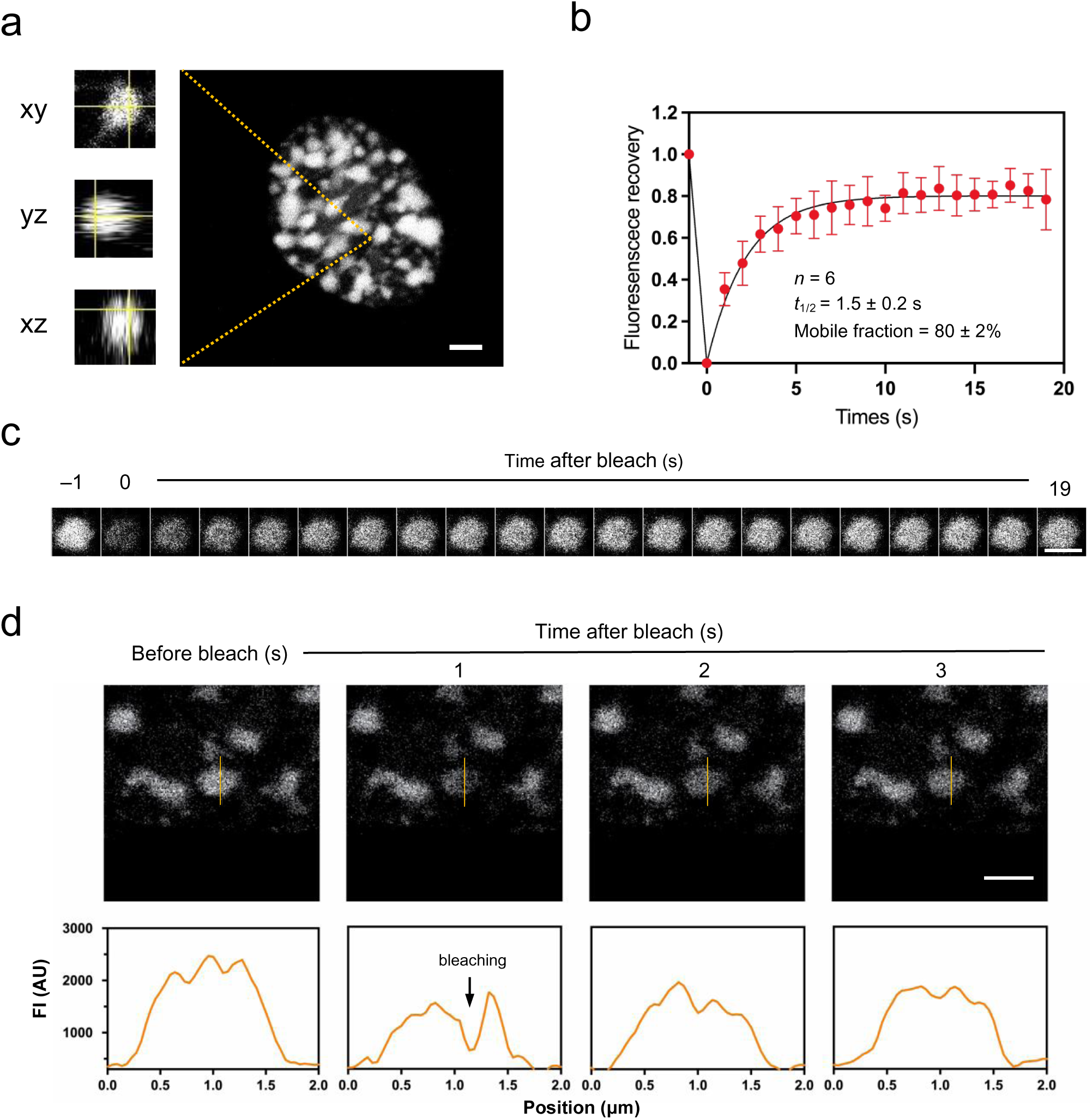
HMGA1a forms condensate foci in the nucleus when overexpressed in IMR90 fibroblasts. **(a)** GFP-HMGA1a proteins nucleate and form spherical, droplet-like foci. **(b)** Full FRAP of HMGA1a foci. Shown is the average signal from 6 condensate foci. HMGA1a foci recover on a timescale of 1.5 s, and the mobile fraction is 80 ± 2%. Data points are mean values; error bars indicate standard deviation. **(c)** Exemplary image series from FRAP experiments as performed in panel b. **(d)** Partial FRAP of HMGA1a foci. The intensity profiles shown in the lower panels correspond to the orange lines of the above images. Scale bars, 2 μm.

To further assess the liquid-like characteristic of HMGA1a condensates, we performed fluorescence recovery after photobleaching (FRAP) experiments on HMGA1a foci. These experiments revealed fast recovery times (average *t*_1/2_ = 1.5 s) with a high mobile fraction (80%) **(Figure 4b)**. This exchange rate (1.5 s) is comparable with, or even faster than, those of many molecules within nuclear biomolecular condensates,^68^ and the mobile fraction is higher than for many heterochromatin proteins, such as HP1a (50%), which has been shown to be able to form phase-separated condensates in *Drosophila* and mammalian cells.^32,33^ These results support the dynamic liquid-like behaviour of GFP-HMGA1a within HMGA condensates.^29,69,70^ Owing to the small size of HMGA1a foci, fluorescence recovery determined by FRAP, as performed here, probes both the entry and exit of EGFP-HMGA1a molecules to the foci from the fluid phase in addition to the mobility of EGFP-HMGA1a in the condensate **(Figure 4c)**. To probe HMGA1a fluidity exclusively within the condensates, we performed partial bleaching experiments **(Figure 4d)**. Fluorescence rapidly recovered (in less than 1 s) from HMGA1a foci. This corroborates that EGFP-HMGA1a foci exhibit liquid-like characteristics.

Importantly, a critical concentration of macromolecules is often needed to trigger prominent phase separation in vivo. Indeed, we found that weaker expression of HMGA1a did abolish the formation of droplet-like foci, or condensates in cells **(Figure S4)**. To investigate this correlation further, we performed experiments using an IMR90 cell population with variable overexpression of HMGA1a, and correlated expression levels of EGFP-tagged HMGA1a with the size and number of resulting condensate structures **(Figure 5a top panels, Figure S5)**. Cells were counterstained with DAPI to provide another mean of staining nuclear condensate structures. This allowed us to compare the nuclear condensate features of cells with low and high levels of HMGA1a. As evident in **Figure 5a and Figure S5**, HMGA1a intensity was variable amongst the cell population, confirming the heterogeneous phenotype in terms of the HMGA1a overexpression. Both DAPI and HMGA1a staining exhibited the formation of droplet-like foci, which were larger and more well-defined in nuclei with stronger HMGA1a overexpression and largely non-existent in cells without HMGA1a overexpression. We quantified the dependency of the condensate structures on HMGA1a intensity across the cell population imaged and observed a positive correlation between the HMGA1a expression level and the average condensate size (normalised by the nucleus size), as well as between the HMGA1a expression level and the number of HMGA1 condensates identified based on HMGA1a staining **(Figure 5b)**.

**Figure 5.**
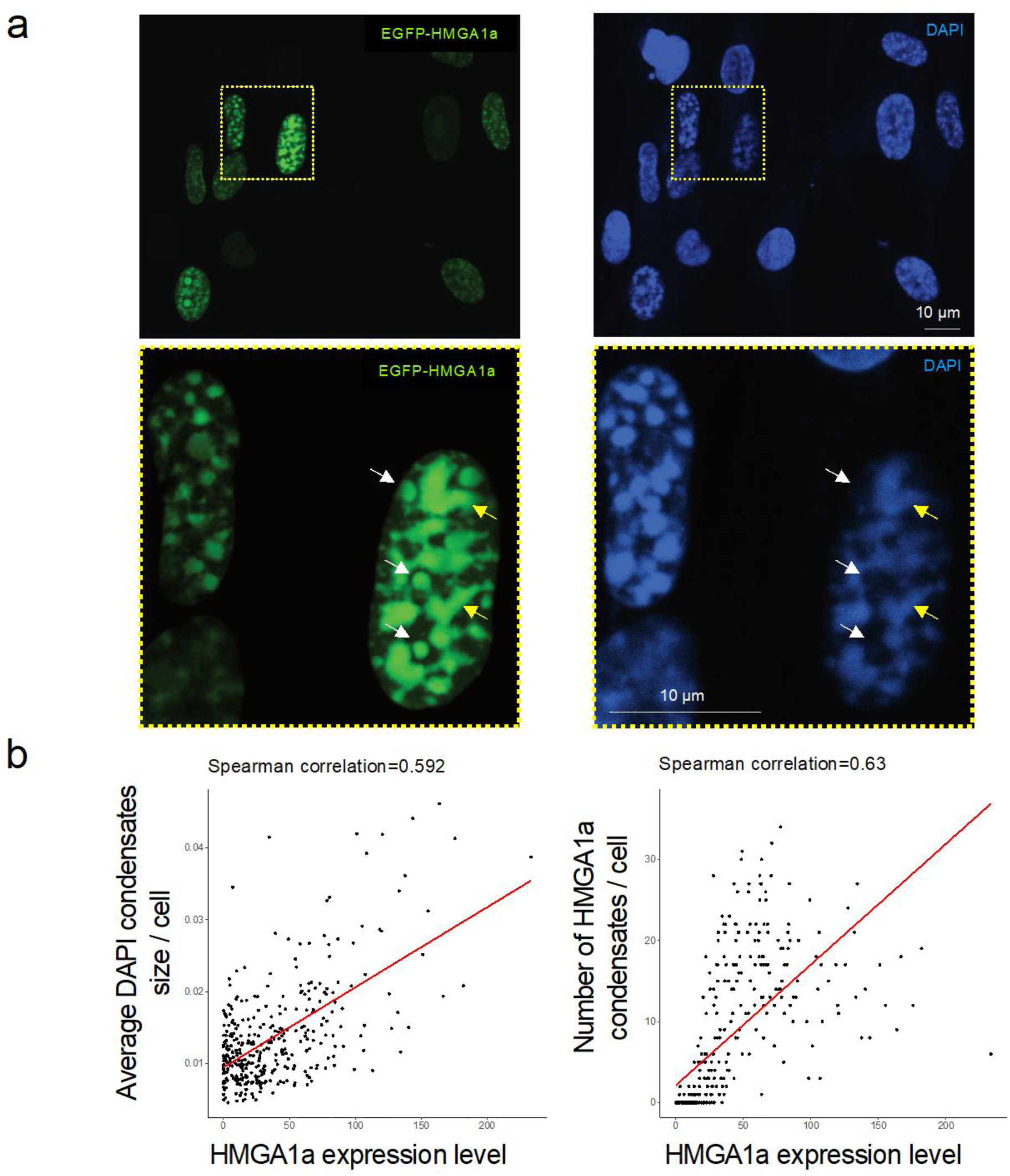
HMGA1a forms condensate foci in the nucleus of IMR90 fibroblasts whose size depend on HMGA1 expression levels. **(a)** Representative confocal images of EGFP-tagged HMGA1a overexpression in IMR90 cells with condensate foci visible inside the nucleus. Cells were counterstained by DAPI to visualize condensate structures. HMGA1a intensity is variable amongst the cell population. The formation of condensates is dependent on HMGA1a expression levels. Three biological replicates were performed. Zoom-ins are shown in the lower panels. Note that at DNA-dilute regions, GFP-HMGA1a forms droplet-like, spherical structure, indicated by white arrows. At DNA-rich regions, three structures appear elongated and are of irregular shape, indicated by yellow arrows. **(b)** Correlation of HMGA1a overexpression with condensate size. Left: HMGA1 expression level per cell nucleus versus average size of condensates per cell nucleus. The signal was normalised by the nucleus size. Right: HMGA1 expression level per cell nucleus versus number of HMGA1 condensates detected in each cell nucleus. Spearman’s correlation coefficient is given as an inset. The trend line is obtained from linear fitting.

Taking a closer look at the HMGA1a condensates in cells **(Figure 5a, lower panels)**, we observed that at DNA-dilute regions GFP-HMGA1a forms well-defined spherical structures, indicated by white arrows. At DNA-rich regions, there structures appear elongated and are of irregular shape, indicated by yellow arrows. We speculate the loss of circularity is a result of more HMGA1a associating with the chromatin polymer, whose inherent elasticity introduces shape constraints.^71^

Interestingly, treatment with 1,6-hexanediol did not suppress the formation of HMGA1a foci in the nucleus **(Figure S6)**, suggesting that hydrophobic interactions are likely not predominant in the formation of HMGA1a condensates. While for many biological systems, hydrophobic interactions are the main driving force of LLPS, other types of interactions, including π–π, cation-π, and electrostatic interactions, have been shown to sustain LLPS; the latter two interactions are relatively unaffected by addition of hexanediol.^72–74^

We also probed LLPS of HMGA1a in a different cell line, by expressing mVenus-fused HMGA1a in the HCT116 human colon cancer cell line. Images obtained via microscopy revealed similar HMGA1a foci to those observed in IMR90 nuclei, as well as rapid FRAP recovery **(Figure S7)**. Again, in HCT116 cells, HMGA1a foci were only observed in the nuclei with high expression of HMGA1a. Notably, fewer HMGA1a foci were observed in HCT116 nuclei than in IMR90 nuclei. This result suggests a possible dependency on cell type for HMGA1a droplet formation, yet there may be additional factors to IMR90 that regulate HMGA1a LLPS depending on the cellular environment.

## Conclusions

We have shown that HMGA1a can undergo phase separation to form liquid condensates in vitro and nucleates into foci that display liquid properties in fibroblasts and cancer cells. Both experimental results and modelling data show that LLPS of HMGA1a is promoted in the presence of DNA. HMGA1a–DNA condensates are possibly stabilized by dominant cation– π and electrostatic interactions. We suggest that the formation of liquid-like domains enriched in HMGA1a in the nucleus is contributed by both homotypic HMGA1a interactions and heterotypic interactions with nucleosomes and DNA. We further speculate that the formation of HMGA1a liquid droplets in cells within the regions depleted of DNA and nucleosomes might be enabled by post-translational modifications of HMGA1a (i.e., phosphorylation). Indeed, an increased residence time of HMGA1a to heterochromatin regions, which would be consistent with its LLPS, has been shown to be correlated with high levels of phosphorylation.^5^

Importantly, other architectural proteins that are also enriched in heterochromatin regions, like the linker histone H1, and the heterochromatin protein HP1, have also been observed to undergo LLPS in vitro and in cells. There are many parallels between the phase behaviour of these proteins and our observations for HMGA1a. In particular, LLPS of H1 is dependent on the presence of DNA or nucleosomes, and LLPS of HP1 is dependent on the phosphorylation of the negatively charged N-terminal region, or the presence of DNA.^30,32,33,75^

The sensitivity of HMGA1a towards the presence of post-translational modifications^61^ is consistent with the functional importance of this protein, and hence, the need for its behaviour to be highly regulated. A hypothesis stemming from our work, is that different post-translational modifications of HMGs can lead to the formation of diverse nuclear condensates that selective recruit or exclude DNA regions, perhaps to control gene function.

## Methods

### Phase separation prediction

LLPS propensity of HMGA1a was modelled using the DeePhase algorithm.^52^ Briefly, the model converted the sequence into an input vector consisting of a number of explicit sequence-specific parameters (sequence length, hydrophobicity, Shannon entropy, the fraction of polar, aromatic and positively charged residues and the fraction of sequence estimated to be part of the low complexity region and intrinsically disordered region) as well as implicit word2vec algorithm-based embeddings. The model had been trained on datasets including sequences with varying level of disorder and varying propensity to undergo LLPS. To estimate the local LLPS-propensity across the HMGA1a sequence, the full sequence was divided into 30 amino acid long fragments and the propensity of each fragment to undergo LLPS was evaluated separately. The result was averaged using a running mean with window size of 7. We note that while the score corresponds to the propensity of specific regions along the sequence to undergo phase separation, globally, regions with low LLPS-propensity can play an essential role in facilitating the phase separation process.

### In vivo condensate imaging and FRAP

EGFP-HMGA1a was stably expressed in IMR90 cells (ATCC) via retroviral gene transfer with either a strong (CMV) or a weaker (LTR) promoter.^15^ mVenus-HMGA1a was stably expressed in HCT116 cells (ATCC) using the *PiggyBac* transposon system.^76^ In vivo HMGA1a condensate foci were visualised by a z-stack imaging mode at a single-cell level using a Leica TCS SP8 confocal microscope. FRAP experiments were performed on in vivo condensates formed by GFP-HMGA1a or mVenus-HMGA1a using the 488 or 514 nm laser line, respectively, using a Leica TCS SP8 confocal microscope. FRAP in cells was performed on a selected point with 100% power (50 ms) and recovery observed at 2% power, 1 s intervals for 20 s. Image analysis was performed with Fiji. Recovery was measured as fluorescence intensity of photobleached area normalised to the intensity of the unbleached area. Immobile fractions were measured as percent fluorescence intensity unrecovered after 20 s.

### Recombinant HMGA1a expression and purification

To produce the full-length recombinant human HMGA1a protein, the pRSET-A expression vector was transformed into the double lon/omp T protease mutant B strain of *E. coli* BL21(DE3)pLysS.^77^ Recombinant Human HMGA1a was expressed in *E. coli* BL21(DE3)pLysS upon IPTG induction and purified in two steps by immobilized metal affinity chromatography (IMAC) and cation exchange chromatography using a HiTrap SP HP column (Cytiva). The buffers used with HiTrap SP HP column were 10 mM Tris (pH 7.4), 300 mM NaCl (low salt) and 10 mM Tris (pH 7.4), 1 M NaCl (high salt). The purity of each recombinant preparation was assessed by SDS-PAGE. Protein concentrations were determined spectrophotometrically employing either a Bio-Rad protein assay kit or using the extinction coefficient ε_220_ = 74,000 L/mol·cm for HMGA1a protein.^78^

### In vitro droplets assays

Manual in-vitro droplet assays were performed by mixing indicated final amounts of protein, double-stranded DNA stocks, and PEG (20k, Sigma) in 50 mM Tris buffer (pH 7.4). The protein stock was labelled with Alexa 647-*N*-Hydroxysuccinimide (Alexa 647-NHS, Thermo Fisher) at sub-stoichiometric ratios yielding a labelling efficiency of <5%. Duplex DNA was prepared from two single-stranded DNA oligonucleotides by thermal annealing. Oligonucleotides were synthesized and labelled by IDT. The sequences were: 5’-CAC AAC TCC GCT GCG TCA GAG CAG-3’ (top strand) and 5’-CTG CTC TGA CGC AGC GGA GTT GTG-3’ (bottom strand); the top strand was labelled with Atto488 at the 5’-end. Phase-separated samples were prepared in tubes and imaged within 1–5 min. Imaging was performed on an inverted fluorescence microscope (OpenFrame, Cairn Research) equipped with a high-sensitivity camera (Prime BSI Express, Photometrics) by placing an aliquot of the sample (1–2 μL) between two coverslips. Samples were imaged using an Olympus 100x NA 1.4 oil-immersion objective. Appropriate filter sets for Atto 488 and Alexa 647 detection were used.

### PhaseScan

Phase diagrams were produced using droplet microfluidics in a similar manner to that described previously,^64^ using polydimethylsiloxane (Corning) devices produced on SU-8 (Microchem) moulds which were fabricated via photolithographic processes.^79–81^ Syringe pumps (neMESYS modules, Cetoni) were used to control flows of input solutions of HMGA1a, 3.6 μM or 0.2 μM Atto 488 labelled duplex DNA, buffer (50 mM Tris (pH 7.4) 120 mM KCl), and PEG 20k (15% *w*/*v*) supplemented with 3 μM Alexa 546 dye (carboxylic acid, ThermoFisher). The protein solution consisted of 70 μM HMGA1a supplemented with 10 μM Alexa647-labelled HMGA1a, HMGA1a was labelled with Alexa 647 dye by incubating the protein in 1:1 molar ratio with Alexa 647-NHS ester for 20 min at room temperature. The aqueous flow rates were configured to vary automatically according to pre-set gradients, with constant total flow rate of 60 μL/h, to scan phase space between nominal concentrations of 3–47 μM and 0.01–2.1 μM for HMGA1a and DNA, respectively. FC-40 oil (containing 1% (*w*/*v*) fluorosurfactant, RAN biotechnologies) was introduced to the device at a constant flow rate of 150 μL/h for microdroplet generation. After generation, microdroplets were incubated on chip for 2.5 min during passage through a flow channel, before being imaged under flow on a custom-built epifluorescence microscope (OpenFrame, Cairn Research) equipped with a 10x air objective, high-sensitivity camera (Kinetix sCMOS, Photometrics) and optical splitter (Multisplit, Cairn Research).

### Simulations

#### HMGA1a model

HMGA1a protein was modelled using the HPS model of Dignon et al.^58^ with a modification introduced by Das et al.^60^ to account for cation–π interactions. In this model each protein residues is modelled via a single bead that has a unique charge, hydrophobicity (i.e., hydropathy score), mass, and van der Waals radius. Within this framework, the energy of the system is computed as the sum of a scaled Lennard-Jones interaction for pairwise contacts, Coulombic Debye–Huckel term for long-range electrostatic interactions, and a standard harmonic potential for bonded interactions. The sequence of HMGA1a was obtained from Uniprot^82^ and was mapped unto a random chain using Pymol software.^83^ In our simulation, each HMGA1a protein was represented as a fully flexible chain.

#### DNA model

Double-stranded DNA was represented via our chemically accurate coarse-grained model for DNA.^22^ This model was parametrized to account for the mechanical and chemical properties of DNA. Particularly, each DNA bp is represented via an ellipsoid of appropriate mass (based on the identity of the bases) and with point charges (2 in total) to account for the charged sugar-phosphate backbone. For this work, we use DNA strands composed of 24 bps; i.e., compatible with DNA linker lengths in chromatin. While the model does capture well the changes in DNA mechanical (bending, twisting etc) properties with sequence, at these short lengths the persistence lengths of different DNA sequences are all comparable. Hence, we used a random DNA sequence (See Supporting Information) for this study.

#### Direct coexistence simulations

To probe LLPS behaviour of HMGA1a and DNA, we use the Direct Coexistence (DC) method.^84–86^ In this approach, the protein-rich and protein-depleted phases are both represented in the same simulation box. For the pure HMGA1a system we used 48 copies of the protein (107 residues each), and for the HMGA1a–DNA mixture 48 copies of HMGA1a and 12 strands of double-stranded DNA (each stand = 24 base pairs). Each system was first prepared in a cubic box. Isotropic *NPT*-ensemble (constant pressure and temperature) simulations were then performed at high pressure (>20 bars; using a Berendsen barostat) and low temperature (temperature regulated via a Langevin thermostat) to produce a high-density slab-like structure. One side of the box was then elongated (ca. 3– 10 times the box cross section) and *NVT*-ensemble simulations were then performed. Each system was simulated for approx. 2–5 microseconds. To assess convergence, the density and the energy of the system were monitored. The presence of the well-defined interface was used to indicate LLPS, while, the lack of such interface is indicative of no LLPS under a given set of conditions. Here, we report the temperature of our systems in terms of the critical temperature of the pure HMGA1a wildtype system (referred to as *T*_c_(wt) ∼200 K). All simulations were performed using the LAMMPS simulation package.^87^

#### Contact map analysis (contact frequency)

At a given temperature, the contact frequency between protein residues (and between protein residues and DNA bps) was measured using the Python MDAnalysis package.^88,89^ Two residues (*i* and *j*) (or a residue and a DNA base pair) were deemed to be in contact if they are within a threshold of *r*_ij_; where *r*_ij_ is the average of their respective molecular diameters.

#### Estimation of DNA valency

At 1.85 *T*_c_(wt), each protein–DNA contact contributes approximately 0.45kT to the interaction energy. Hence, 3 of these contacts are required to make a sizable contribution to the overall interaction energy. Accordingly, we imposed the condition that to be “in contact” at least 3 of these contacts must exist between a DNA strand and a protein chain (i.e., a contribution of +1 to the valency of DNA valency). Using this condition with the contact analysis approach (explained above), we find that each DNA strand (24 bps) recruits on average 4.49 proteins (ca. 4–5 proteins). Hence, we conclude that 12 bps can successfully bridge 2 HMGA1a proteins. The analysis was performed using the Python MDAnalysis package.^88,89^

### Cell nuclei and condensate structure detection

We used the StarDist package^90,91^ to detect nuclear contours from images with DAPI staining in order to define cell nuclei and also to detect condensate structures in images of DAPI and HMGA1 staining. The condensate structures were identified by iterating and performing segmentation on each individual cell nucleus. The sizes of the object identified, as well as average HMGA1 intensity per nucleus were quantified using the scikit-image Python module^92^.

## Acknowledgements

This project has received funding from the European Research Council (ERC) under the European Union’s Horizon 2020 research and innovation programme (grant agreement No 803326) and Cancer Research UK Cambridge Institute Core Grant (C9545/A29580). H.Z. received funding from China Scholarship Council (CSC). J.A.J. is a Research Fellow at King’s College. R.C.-G. is an Advanced Fellow from the Winton Programme for the Physics of Sustainability. J.R.E acknowledges funding from Oppenheimer and Roger Ekins fellowships. G.K. acknowledges funding from the Herchel Smith Funds and the Wolfson College Junior Research Fellowship. K.L.S. is supported by the Schmidt Science Fellowship programme in partnership with the Rhodes Trust. This work has been performed using resources provided by the Cambridge Tier-2 system operated by the University of Cambridge Research Computing Service (http://www.hpc.cam.ac.uk) funded by EPSRC Tier-2 capital grant EP/P020259/1.

## References

1. Maeshima, K., Tamura, S., Hansen, J. C. & Itoh, Y. Fluid-like chromatin: Toward understanding the real chromatin organization present in the cell. Curr. Opin. Cell Biol. 64, 77–89 (2020).

2. Sanulli, S. et al. HP1 reshapes nucleosome core to promote phase separation of heterochromatin. Nature 575, 390–394 (2019).

3. Ozturk, N., Singh, I., Mehta, A., Braun, T. & Barreto, G. HMGA proteins as modulators of chromatin structure during transcriptional activation. Front. cell Dev. Biol. 2, 5 (2014).

4. Postnikov, Y.V & Bustin, M. Functional interplay between histone H1 and HMG proteins in chromatin. Biochim. Biophys. Acta 1859, 462–467 (2016).

5. Harrer, M., Lührs, H., Bustin, M., Scheer, U. & Hock, R. Dynamic interaction of HMGA1a proteins with chromatin. J. Cell Sci. 117, 3459–3471 (2004).

6. R, H., T, F., T, U. & M, B. HMG chromosomal proteins in development and disease. Trends Cell Biol. 17, 72–79 (2007).

7. Pallante, P., Sepe, R., Puca, F. & Fusco, A. High Mobility Group A Proteins as Tumor Markers. Front. Med. 0, 15 (2015).

8. A, F. & M, F. Roles of HMGA proteins in cancer. Nat. Rev. Cancer 7, 899–910 (2007).

9. Li, O., Vasudevan, D., Davey, C. A. & Dröge, P. High-level expression of DNA architectural factor HMGA2 and its association with nucleosomes in human embryonic stem cells. genesis 44, 523–529 (2006).

10. Brocher, J., Vogel, B. & Hock, R. HMGA1 down-regulation is crucial for chromatin composition and a gene expression profile permitting myogenic differentiation. BMC Cell Biol. 11, 64 (2010).

11. Lanahan, A., Williams, J. B., Sanders, L. K. & Nathans, D. Growth factor-induced delayed early response genes. Mol. Cell. Biol. 12, 3919–3929 (1992).

12. Zanin, R. et al. HMGA1 promotes breast cancer angiogenesis supporting the stability, nuclear localization and transcriptional activity of FOXM1. J. Exp. Clin. Cancer Res. 38, 313 (2019).

13. Fu, F. et al. HMGA1 exacerbates tumor growth through regulating the cell cycle and accelerates migration/invasion via targeting miR-221/222 in cervical cancer. Cell Death Dis. 9, 594 (2018).

14. Chen, J.-H., Hales, C. N. & Ozanne, S. E. DNA damage, cellular senescence and organismal ageing: causal or correlative? Nucleic Acids Res. 35, 7417–7428 (2007).

15. Narita, M. et al. A Novel Role for High-Mobility Group A Proteins in Cellular Senescence and Heterochromatin Formation. Cell 126, 503–514 (2006).

16. Tremethick, D. J. Higher-Order Structures of Chromatin: The Elusive 30 nm Fiber. Cell 128, 651–654 (2007).

17. Maeshima, K., Hihara, S. & Eltsov, M. Chromatin structure: Does the 30-nm fibre exist in vivo? Curr. Opin. Cell Biol. 22, 291–297 (2010).

18. Collepardo-Guevara, R. & Schlick, T. Chromatin fiber polymorphism triggered by variations of DNA linker lengths. Proc. Natl. Acad. Sci. U. S. A. 111, 8061–8066 (2014).

19. Krietenstein, N. & Rando, O. J. Mesoscale organization of the chromatin fiber. Curr. Opin. Genet. Dev. 61, 32–36 (2020).

20. Ricci, M. A., Manzo, C., García-Parajo, M. F., Lakadamyali, M. & Cosma, M. P. Chromatin fibers are formed by heterogeneous groups of nucleosomes in vivo. Cell 160, 1145–1158 (2015).

21. Itoh, Y., Woods, E. J., Minami, K., Maeshima, K. & Collepardo-Guevara, R. Liquid-like chromatin in the cell: What can we learn from imaging and computational modeling? Curr. Opin. Struct. Biol. 71, 123–135 (2021).

22. Farr, S. E., Woods, E. J., Joseph, J. A., Garaizar, A. & Collepardo-Guevara, R. Nucleosome plasticity is a critical element of chromatin liquid–liquid phase separation and multivalent nucleosome interactions. bioRxiv 2020.11.23.391599 (2020) doi:10.1101/2020.11.23.391599.

23. Brangwynne, C. P. et al. Germline P Granules Are Liquid Droplets That Localize by Controlled Dissolution/Condensation. Science (80-.). 324, 1729–1732 (2009).

24. Lin, Y., Protter, D. S. W. W., Rosen, M. K. & Parker, R. Formation and Maturation of Phase-Separated Liquid Droplets by RNA-Binding Proteins. Mol. Cell 60, 208–219 (2015).

25. Brangwynne, C. P., Tompa, P. & Pappu, R. V. Polymer physics of intracellular phase transitions. Nat. Phys. 11, 899–904 (2015).

26. Hyman, A. A., Weber, C. A. & Jülicher, F. Liquid-Liquid Phase Separation in Biology. Annu. Rev. Cell Dev. Biol. 30, 39–58 (2014).

27. Shin, Y. & Brangwynne, C. P. Liquid phase condensation in cell physiology and disease. Science (80-.). 357, (2017).

28. Ditlev, J. A., Case, L. B. & Rosen, M. K. Who’s in and who’s out—compositional control of biomolecular condensates. J. Mol. Biol. (2018).

29. Alberti, S., Gladfelter, A. & Mittag, T. Considerations and Challenges in Studying Liquid-Liquid Phase Separation and Biomolecular Condensates. Cell 176, 419–434 (2019).

30. Gibson, B. A. et al. Organization of Chromatin by Intrinsic and Regulated Phase Separation. Cell 179, 470–484 (2019).

31. Strom, A. R. & Brangwynne, C. P. The liquid nucleome – phase transitions in the nucleus at a glance. J. Cell Sci. 132, (2019).

32. Larson, A. G. et al. Liquid droplet formation by HP1α suggests a role for phase separation in heterochromatin. Nature 547, 236–240 (2017).

33. Strom, A. R. et al. Phase separation drives heterochromatin domain formation. Nature 547, 241 (2017).

34. Wang, J. et al. A Molecular Grammar Governing the Driving Forces for Phase Separation of Prion-like RNA Binding Proteins. Cell 174, 688–699 (2018).

35. Martin, E. W. et al. Valence and patterning of aromatic residues determine the phase behavior of prion-like domains. Science (80-.). 367, 694–699 (2020).

36. Espinosa, J. R. et al. Liquid network connectivity regulates the stability and composition of biomolecular condensates with many components. Proc. Natl. Acad. Sci. U. S. A. 117, 13238–13247 (2020).

37. Berry, J. et al. RNA transcription modulates phase transition-driven nuclear body assembly. Proc. Natl. Acad. Sci. U. S. A. 112, E5237–E5245 (2015).

38. Saha, S. et al. Polar Positioning of Phase-Separated Liquid Compartments in Cells Regulated by an mRNA Competition Mechanism. Cell 166, 1572–1584 (2016).

39. Maharana, S. et al. RNA buffers the phase separation behavior of prion-like RNA binding proteins. Science (80-.). 360, 918–921 (2018).

40. Langdon, E. M. & Gladfelter, A. S. A New Lens for RNA Localization: Liquid-Liquid Phase Separation. Annu. Rev. Microbiol. 72, 255–271 (2018).

41. Rhine, K., Vidaurre, V. & Myong, S. RNA Droplets. Annu. Rev. Biophys. 49, 247–265 (2020).

42. Joseph, J. A. et al. Thermodynamics and kinetics of phase separation of protein-RNA mixtures by a minimal model. Biophys. J. (2021) doi:https://doi.org/10.1016/j.bpj.2021.01.031.

43. Banani, S. F. et al. Compositional Control of Phase-Separated Cellular Bodies. Cell 166, 651–663 (2016).

44. Banani, S. F., Lee, H. O., Hyman, A. A. & Rosen, M. K. Biomolecular condensates: organizers of cellular biochemistry. Nat. Rev. Mol. Cell Biol. 18, 285–298 (2017).

45. Xu, Y. et al. Liquid–Liquid Phase Separated Systems from Reversible Gel–Sol Transition of Protein Microgels. Adv. Mater. 2008670 (2021).

46. Mitrea, D. M. et al. Nucleophosmin integrates within the nucleolus via multi-modal interactions with proteins displaying R-rich linear motifs and rRNA. Elife 5, e13571 (2016).

47. Falahati, H., Pelham-Webb, B., Blythe, S. & Wieschaus, E. Nucleation by rRNA Dictates the Precision of Nucleolus Assembly. Curr. Biol. 26, 277–285 (2016).

48. Fei, J. et al. Quantitative analysis of multilayer organization of proteins and RNA in nuclear speckles at super resolution. J. Cell Sci. 130, 4180–4192 (2017).

49. Nott, T. J. et al. Phase Transition of a Disordered Nuage Protein Generates Environmentally Responsive Membraneless Organelles. Mol. Cell 57, 936–947 (2015).

50. Shakya, A., Park, S., Rana, N. & King, J. T. Liquid-Liquid Phase Separation of Histone Proteins in Cells: Role in Chromatin Organization. Biophys. J. 118, 753–764 (2020).

51. Martin, E. W. & Mittag, T. Relationship of Sequence and Phase Separation in Protein Low-Complexity Regions. Biochemistry vol. 57 2478–2487 (2018).

52. Saar, K. L. et al. Learning the molecular grammar of protein condensates from sequence determinants and embeddings. Proc. Natl. Acad. Sci. 118, e2019053118 (2021).

53. Kato, M. et al. Cell-free formation of RNA granules: Low complexity sequence domains form dynamic fibers within hydrogels. Cell 149, 753–767 (2012).

54. Bremer, A. et al. Deciphering how naturally occurring sequence features impact the phase behaviors of disordered prion-like domains. bioRxiv 2021.01.01.425046 (2021) doi:10.1101/2021.01.01.425046.

55. Patel, A. et al. A Liquid-to-Solid Phase Transition of the ALS Protein FUS Accelerated by Disease Mutation. Cell 162, 1066–1077 (2015).

56. Elbaum-Garfinkle, S. et al. The disordered P granule protein LAF-1 drives phase separation into droplets with tunable viscosity and dynamics. Proc. Natl. Acad. Sci. U. S. A. 112, 7189–7194 (2015).

57. Lin, Y., Currie, S. L. & Rosen, M. K. Intrinsically disordered sequences enable modulation of protein phase separation through distributed tyrosine motifs. J. Biol. Chem. 292, 19110–19120 (2017).

58. Dignon, G. L., Zheng, W., Kim, Y. C., Best, R. B. & Mittal, J. Sequence determinants of protein phase behavior from a coarse-grained model. PLOS Comput. Biol. 14, e1005941 (2018).

59. Li, P. et al. Phase transitions in the assembly of multivalent signalling proteins. Nature 483, 336–340 (2012).

60. Das, S., Lin, Y.-H., Vernon, R. M., Forman-Kay, J. D. & Chan, H. S. Comparative Roles of Charge, Pi, and Hydrophobic Interactions in Sequence-Dependent Phase Separation of Intrinsically Disordered Proteins. arXiv 2005.06712 (2020).

61. Zhang, Q. & Wang, Y. High mobility group proteins and their post-translational modifications. Biochim. Biophys. Acta 1784, 1159–1166 (2008).

62. Kohl, B., Zhong, X., Herrmann, C. & Stoll, R. Phosphorylation orchestrates the structural ensemble of the intrinsically disordered protein HMGA1a and modulates its DNA binding to the NFκB promoter. Nucleic Acids Res. 47, 11906–11920 (2019).

63. Piekielko, A. et al. Distinct Organization of DNA Complexes of Various HMGI/Y Family Proteins and Their Modulation upon Mitotic Phosphorylation*. J. Biol. Chem. 276, 1984–1992 (2001).

64. Arter, W. E. et al. Rapid characterisation of protein phase behaviour using droplet microfluidics. bioRxiv 2020.06.04.132308 (2020) doi:10.1101/2020.06.04.132308.

65. Murakami, T. et al. ALS/FTD Mutation-Induced Phase Transition of FUS Liquid Droplets and Reversible Hydrogels into Irreversible Hydrogels Impairs RNP Granule Function. Neuron 88, 678–690 (2015).

66. Schmidt, H. B. & Gorlich, D. Transport Selectivity of Nuclear Pores, Phase Separation, and Membraneless Organelles. Trends Biochem. Sci. 41, 46–61 (2016).

67. Linsenmeier, M. et al. Dynamics of synthetic membraneless organelles in microfluidic droplets. Angew. Chemie Int. Ed. 1–7 (2019) doi:10.1002/anie.201907278.

68. Courchaine, E. M., Lu, A. & Neugebauer, K. M. Droplet organelles? EMBO J. 35, 1603–1612 (2016).

69. Fujioka, Y. et al. Phase separation organizes the site of autophagosome formation. Nature 578, 301–305 (2020).

70. Wei, M.-T. et al. Phase behaviour of disordered proteins underlying low density and high permeability of liquid organelles. Nat. Chem. 9, 1118 (2017).

71. Vasquez, P. A. et al. Entropy gives rise to topologically associating domains. Nucleic Acids Res. 44, 5540–5549 (2016).

72. Krainer, G. et al. Reentrant liquid condensate phase of proteins is stabilized by hydrophobic and non-ionic interactions. Nat. Commun. 2021 121 12, 1–14 (2021).

73. Qamar, S. et al. FUS Phase Separation Is Modulated by a Molecular Chaperone and Methylation of Arginine Cation-π Interactions. Cell 173, 720-734.e15 (2018).

74. Vernon, R. M. C. et al. Pi-Pi contacts are an overlooked protein feature relevant to phase separation. Elife 7, (2018).

75. A, S., S, P., N, R. & JT, K. Liquid-Liquid Phase Separation of Histone Proteins in Cells: Role in Chromatin Organization. Biophys. J. 118, 753–764 (2020).

76. J, C. & A, B. Generation of an inducible and optimized piggyBac transposon system. Nucleic Acids Res. 35, (2007).

77. Reeves, R. & Nissen, M. S. B. T.-M.in E. Purification and assays for high mobility group HMG-I(Y) protein function. in Chromatin vol. 304 155–188 (Academic Press, 1999).

78. Himes, S. R. et al. The Role of High-Mobility Group I(Y) Proteins in Expression of IL-2 and T Cell Proliferation. J. Immunol. 164, 3157 LP – 3168 (2000).

79. Mazutis, L. et al. Single-Cell Analysis and Sorting Using Droplet-Based Microfluidics. Nat. Protoc. 8, 870–891 (2013).

80. Arter, W. E. et al. Digital Sensing and Molecular Computation by an Enzyme-Free DNA Circuit. ACS Nano (2020) doi:10.1021/acsnano.0c00628.

81. McDonald, J. C. et al. Fabrication of Microfluidic Systems in Poly(dimethylsiloxane). Electrophoresis 21, 27–40 (2000).

82. Consortium, U. UniProt: the universal protein knowledgebase in 2021. Nucleic Acids Res. 49, D480–D489 (2021).

83. The PyMOL molecular graphics system, version 1.8. (2015).

84. Ladd, A. J. C. & Woodcock, L. V. Triple-point coexistence properties of the lennard-jones system. Chem. Phys. Lett. 51, 155–159 (1977).

85. García Fernández, R., Abascal, J. L. F. & Vega, C. The melting point of ice Ih for common water models calculated from direct coexistence of the solid-liquid interface. J. Chem. Phys. 124, 144506 (2006).

86. Espinosa, J. R., Sanz, E., Valeriani, C. & Vega, C. On fluid-solid direct coexistence simulations: The pseudo-hard sphere model. J. Chem. Phys. 139, 144502 (2013).

87. Plimpton, S. Fast parallel algorithms for short-range molecular dynamics. J. Comput. Phys. 117, 1–19 (1995).

88. Michaud-Agrawal, N., Denning, E. J., Woolf, T. B. & Beckstein, O. MDAnalysis: A toolkit for the analysis of molecular dynamics simulations. J. Comput. Chem. 32, 2319–2327 (2011).

89. Gowers, R. J. et al. MDAnalysis: a Python package for the rapid analysis of molecular dynamics simulations. (2019).

90. Schmidt, U., Weigert, M., Broaddus, C. & Myers, G. Cell Detection with Star-Convex Polygons. Lect. Notes Comput. Sci. (including Subser. Lect. Notes Artif. Intell. Lect. Notes Bioinformatics) 11071 LNCS, 265–273 (2018).

91. Weigert, M., Schmidt, U., Haase, R., Sugawara, K. & Myers, G. Star-convex polyhedra for 3D object detection and segmentation in microscopy. Proc. - 2020 IEEE Winter Conf. Appl. Comput. Vision, WACV 2020 3655–3662 (2020) doi:10.1109/WACV45572.2020.9093435.

92. Van Der Walt, S. et al. Scikit-image: Image processing in python. PeerJ 2014, e453 (2014).

