## Supplementary material for "The chromatin regulator HMGA1a undergoes phase separation in the nucleus": SI

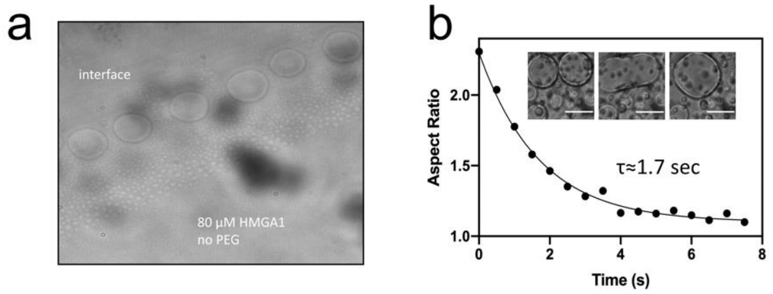


**Figure S1. HMGA1a phase separation in the absence of PEG and fusion propensity of HMGA1a condensates. (a)** In the absence of PEG condensate formation is only observed at the air-water interface. **(b)** Fusion propensity of HMGA1a condensates formed with 5% PEG (20k).

### Supplementary Results

To further characterise the material properties of HMGA1a condensates, we set out to determine their surface tension and viscosity. To this end, PEG and HMGA1a were mixed to trigger phase separation, yielding final concentrations of 10 μM and 5 % (*w*/*w*), respectively, and fusion events were recorded. Fusing HMGA droplet aspect ratios (A.R.) were plotted over time ($t$) **(Figure S1)** and fitted by using least square minimization to A.R.= (A.R.($t_0$) – P) × $e^{-t/\tau}$ + P, where $\tau$ is relaxation time constant, A.R.($t_0$) is the aspect ratio at time $t = 0$, and $P$ the plateau reached at infinite time. The relaxation time constant ($\tau$) was found to be approximately 1.7 s. The size of droplets ($l$) were plotted against $\tau$ to determine the capillary velocity from the slope, the inverse of which is the inverse capillary velocity, $\tau/L = \eta/\gamma$, where L is the condensate diameter, $\eta$ is the viscosity and $\gamma$ the surface tension. We found $\tau/L = \eta/\gamma$ to be 0.0830 s/μm, which is much less viscous than the nucleolus (46 s/μm) and P granules (0.12 s/μm).[55,90] Assuming a typical length scale of $\xi \approx 10$ nm for HMGA1 proteins according to a previous study,[90] this yields $\gamma \approx k_B T / \xi^2 \approx 4 \times 10^{-5}$ K/m$^2$. Knowing the capillary velocity and the interfacial tension, the dynamic viscosity $\eta$ is estimated to be 3.3 Pa s, which is 3300 times greater than that of water, but orders of magnitude less than that of nucleoli (~2000 Pa s).

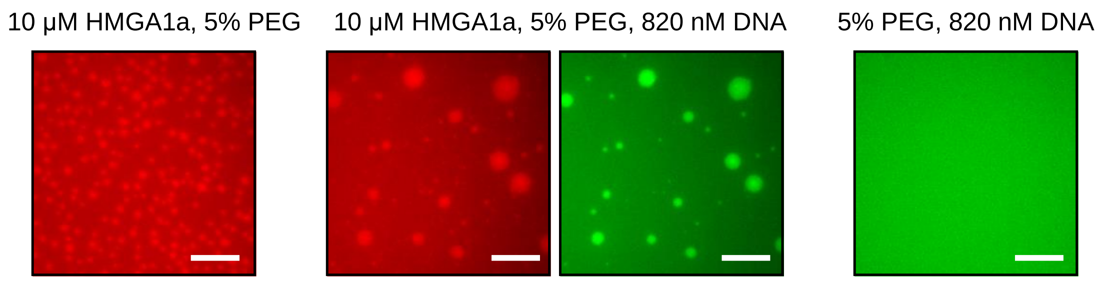


**Figure S2. A control experiment without protein, but only DNA present, did not result in condensate formation.** Scale bar: 10 μm.

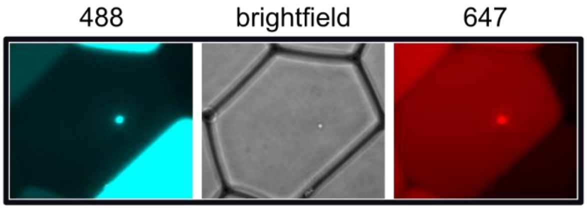


**Figure S3. HMGA1a condensates merge into a single condensate within micro-droplets indicating their liquid-like character.**

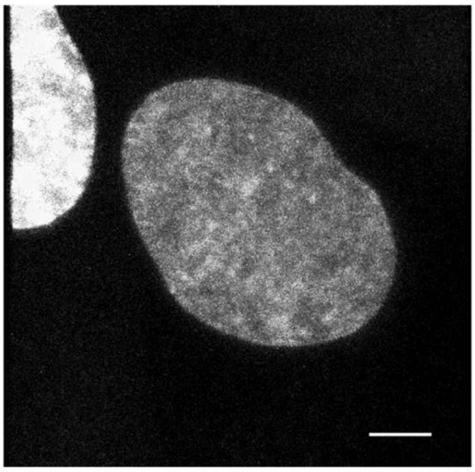

**Figure S4. GFP-HMGA1a under low overexpression levels does not form condensates in IMR90 cells.** Scale bar: 5 μm.

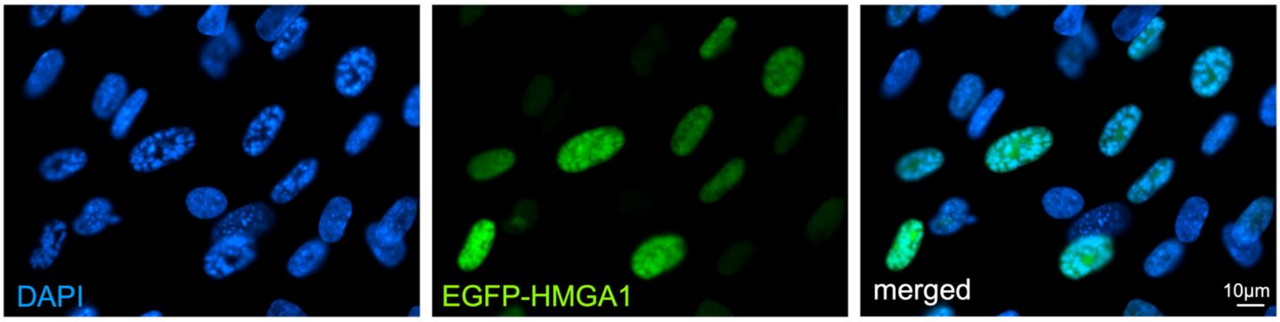


**Figure S5. Representative image of three biological replicates of the EGFP tagged HMGA1 overexpression in IMR90 cells and counterstained by DAPI.**

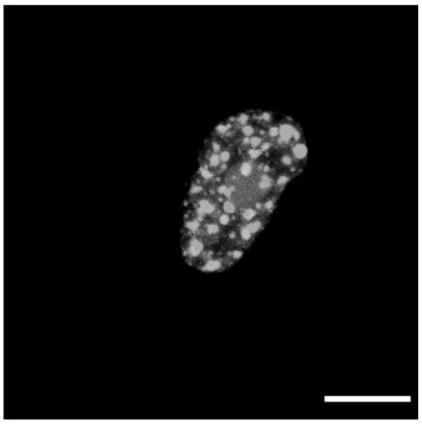

**Figure S6. Hexanediol treatment of GFP-HMGA1a overexpressing cells has no effect on condensate foci.** Scale bar: 5 µm

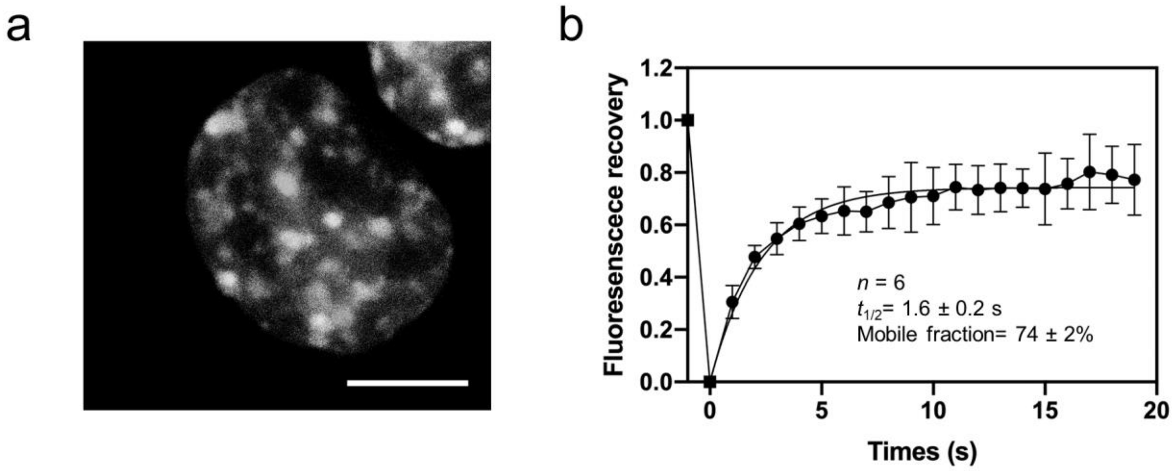


**Figure S7. HMGA1a forms condensate foci in the nucleus when overexpressed in HCT116 cells. (a)**

mVenus-HMGA1a proteins nucleate and form spherical, droplet-like foci. **(b)** Full FRAP of HMGA1a foci. Shown is the average signal from 6 condensate foci. HMGA1 foci recover on a timescale of 1.6 ± 0.2 s, and the mobile fraction is 74 ± 2%.
